# Efficiently Constructing Complete Genomes with CycloneSEQ to Fill Gaps in Bacterial Draft Assemblies

**DOI:** 10.1101/2024.09.05.611410

**Authors:** Hewei Liang, Mengmeng Wang, Tongyuan Hu, Haoyu Wang, Wenxin He, Yanmei Ju, Ruijin Guo, Junyi Chen, Fei Guo, Tao Zeng, Yuliang Dong, Bo Wang, Chuanyu Liu, Xin Jin, Wenwei Zhang, Yuanqiang Zou, Xun Xu, Liang Xiao

**Author notes:** Corresponding author: Yuanqiang Zou, Xun Xu, Liang Xiao.

## Abstract

**Background:** Current microbial sequencing relies on short-read platforms like Illumina and DNBSEQ, favored for their low cost and high accuracy. However, these methods often produce fragmented draft genomes, hindering comprehensive bacterial function analysis. CycloneSEQ, a novel long-read sequencing platform developed by BGI-Research, its sequencing performance and assembly improvements has been evaluated.

**Findings:** Using CycloneSEQ long-read sequencing, the type strain produced long reads with an average length of 11.6 kbp and an average quality score of 14.4. After hybrid assembly with short reads data, the assembled genome exhibited an error rate of only 0.04 mismatches and 0.08 indels per 100 kbp compared to the reference genome. This method was validated across 9 diverse species, successfully assembling complete circular genomes. Hybrid assembly significantly enhances genome completeness by using long reads to fill gaps and accurately assemble multi-copy rRNA genes, which unable be achieved by short reads solely. Through data subsampling, we found that over 500 Mbp of short-read data combined with 100 Mbp of long-read data can result in a high-quality circular assembly. Additionally, using CycloneSEQ long reads effectively improves the assembly of circular complete genomes from mixed microbial communities.

**Conclusions:** CycloneSEQ’s read length is sufficient for circular bacterial genomes, but its base quality needs improvement. Integrating DNBSEQ short reads improved accuracy, resulting in complete and accurate assemblies. This efficient approach can be widely applied in microbial sequencing.

## Background

Current microbial sequencing is primarily based on short-read sequencing technologies [1], including mainstream platforms such as Illumina and DNBSEQ, for both isolated genome and metagenomic studies [2-4]. Short reads are favored for their low cost and high accuracy [5]. However, assemblies based on short reads typically results in a draft genome [6], which are presented as several to several hundred contigs. This fragmented assembly hinders the comprehensive understanding of bacterial functions [7, 8]. In this decade, short-read sequencing has been widely applied in the microbial field, contributing to a large number of diverse draft genomes [2-4]. However, it remains challenging to close the gaps between contigs in these draft genomes using only short reads.

Long-read sequencing, has been developed for nearly two decades [9]. By using long-read assembly, or a hybrid assembly combining long reads with short reads, the genome completeness and proportion of complete circular genomes assembled can be significantly increased [10]. Despite this, the cost of long-read sequencing remains significantly higher than that of short reads [11]. It limits the widespread application of long-read sequencing, especially in large-scale datasets. Currently, there are only a few self-produced large-scale datasets, such as the NCTC3000 [12] and the *Actinomycete* genomes dataset [13]. Complete bacterial genomes can provide comprehensive insights into genomic structures, promote the identification of novel genes, and enhance our understanding of microbial evolution [14].

CycloneSEQ is a long-read sequencing platform developed by BGI-Research, which utilizes nanopore to perform long-read sequencing[15]. It has demonstrated excellent performance in sequencing the *Escherichia coli* genome. However, its performance in sequencing diverse microbial genomes has not yet been systematically evaluated This study focuses on assessing the performance of CycloneSEQ in microbial sequencing and the improvements in genome assembly achieved using CycloneSEQ long-read. By integrating short reads from DNBSEQ with long reads from CycloneSEQ in a hybrid assembly, we validate the effectiveness of this approach in assembling complete circular genomes. And the complete genome will enhance our understanding of bacterial functional gene coding. Additionally, we are interested in the data volume required for assembly. By performing random subsampling of the long-read and short-read sequencing data, we test the assembly performance within data ranges of 100 Mbp, 200 Mbp, 500 Mbp, and 1000 Mbp, providing insights into the success rate of achieving complete assemblies and their accuracy at different data volumes.

## Results

### Sequencing and genome assembly for a type strain

In order to evaluate the accuracy of CycloneSEQ sequencing and the quality of the assembled genome, we cultured a type strain *Akkermansia muciniphila* ATCC BAA-835 and performed CycloneSEQ and DNBSEQ sequencing on its extracted DNA. We have obtained high-depth sequencing data with both short-read and long-read sequencing exceeding 1000x coverage, including a total of 12.07 Gbp of long-reads with an average length of 11,659.2 bp (**Fig. 1A**), and paired short-read with a total of 4.10 Gbp and an average length of 99.9 bp (**Supplementary Table S1**). The average quality of the long reads was 14.4 (**Fig. 1B**), which improved to 14.9 after quality control by selecting reads with a quality value greater than Q10. The paired short reads had an average quality of 35.8 and 34.9, respectively (**Supplementary Table S1**). Furthermore, the use of different sequencing reads and assembly methods can affect the final assembly results. We chose the widely-used software Unicycler [16] (which relies on SPAdes [17] for short-read assembly) and Flye [18] to perform assemblies using only short reads, only long reads, and a hybrid of both.

**Figure 1.**
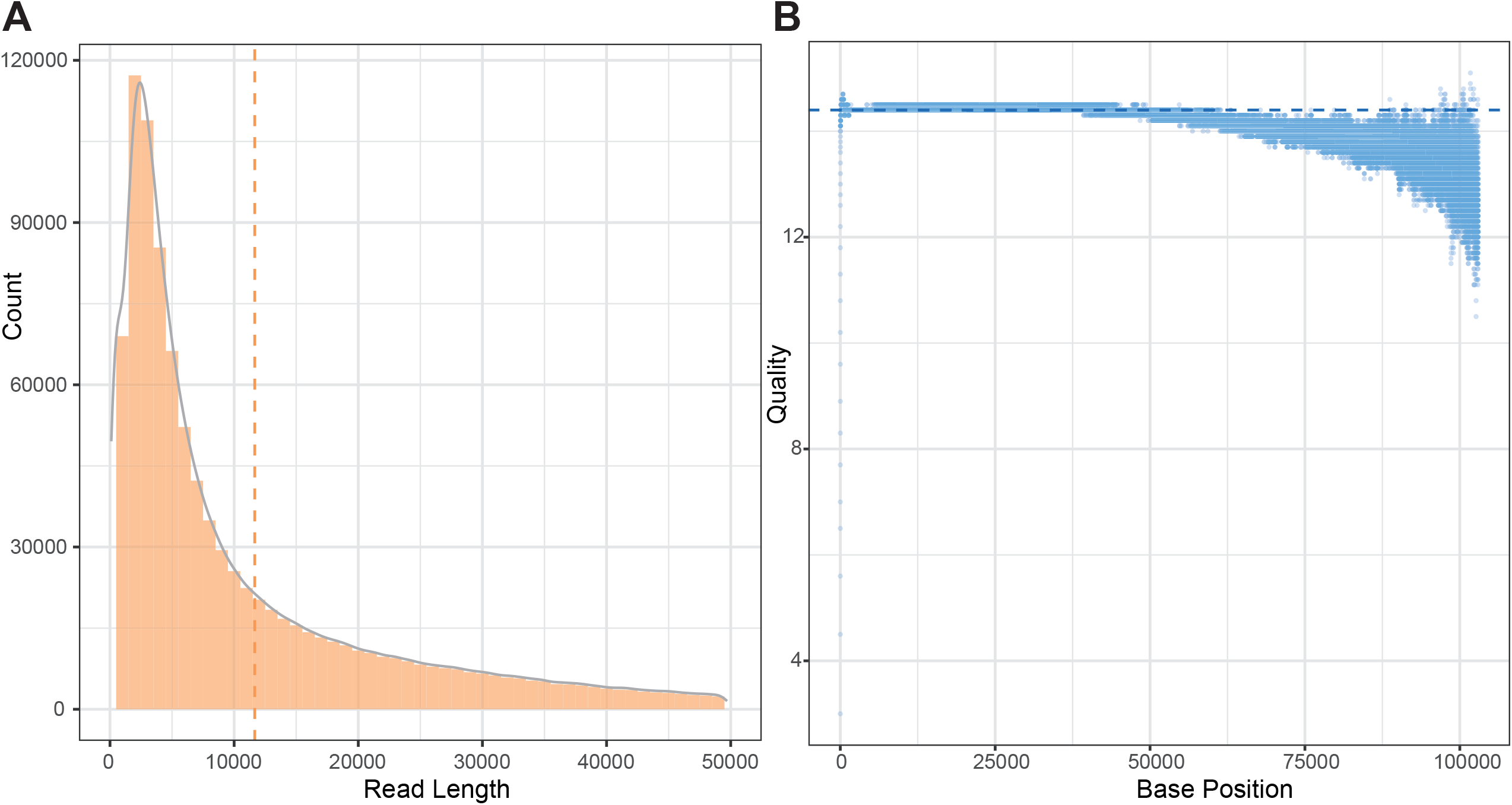
Raw data information from the sequencing. **A**. The bar plot denotes the count of reads in different length ranges, and the curve line denotes the density of read lengths. **B**. The quality of each base position in each read.

Both long-read and hybrid assemblies resulted in a single circular genome, while the short-read assembly resulted in 46 contigs (**Supplementary Fig. 1A**). The GC content was sensitively affected by the completeness and accuracy of the assembly, with the short-read assembly being 55.74%, the long-read assembly being 55.75%, and the hybrid assembly being consistent with the reference at 55.76% (**Supplementary Fig. 1B**). In terms of total length, the hybrid assembly’s length of 2,664,100 was closest to the reference’s 2,664,102, while the long-read assembly was 2,661,711 and the short-read assembly was 2,635,075 (**Supplementary Fig. 1C**), indicating that the short-read assembly had much more fragmentary gaps.

As for the error rate, the short reads achieved a quality of Q35. Thus, short-read assembly exhibited only 0.04 mismatches and 0.08 indels per 100 kbp (**Supplementary Fig. 2**). The hybrid assembly, which was based on the short-read assembly, had 0.08 mismatches and 0.15 indels per 100 kbp. By contrast, the long-read assembly’s error rate was several hundredfold higher, with 13.53 mismatches and 127.49 indels per 100 kbp (**Supplementary Fig. 2**). Such a high error rate could badly affect subsequent analysis of the assembly. Overall, we consider hybrid assembly to be the optimal assembly method.

### Hybrid assembly enhances genome accuracy and completeness

To evaluate the performance of CycloneSEQ and DNBSEQ on actual samples, we collected 10 strains from 9 diverse species for sequencing. The long-read sequencing data for these 10 strains had average quality scores ranging from 12.3 to 15.5 (**Supplementary Table S2**). We then assembled the data using only short reads, only long reads, and a hybrid of both. The hybrid assemblies consistently resulted in circular genomes (**Fig. 2**), including potential small circular genomes from bacteriophages or plasmids. With long-read assemblies, 8 out of 10 strains were successfully assembled into circular genomes, whereas short-read assemblies did not result in circular genomes. When compared to the hybrid assembly genomes, the long-read assemblies exhibited more than a hundred indels and mismatches per 100 kbp, while the short-read assemblies showed almost no variation, with fewer than 3 indels and mismatches per 100kbp (**Supplementary Fig. 3**). These findings are similar to those from type strain analyses, where long-read assemblies tend to be more error-bases, and short-read assemblies are fragmented.

**Figure 2.**
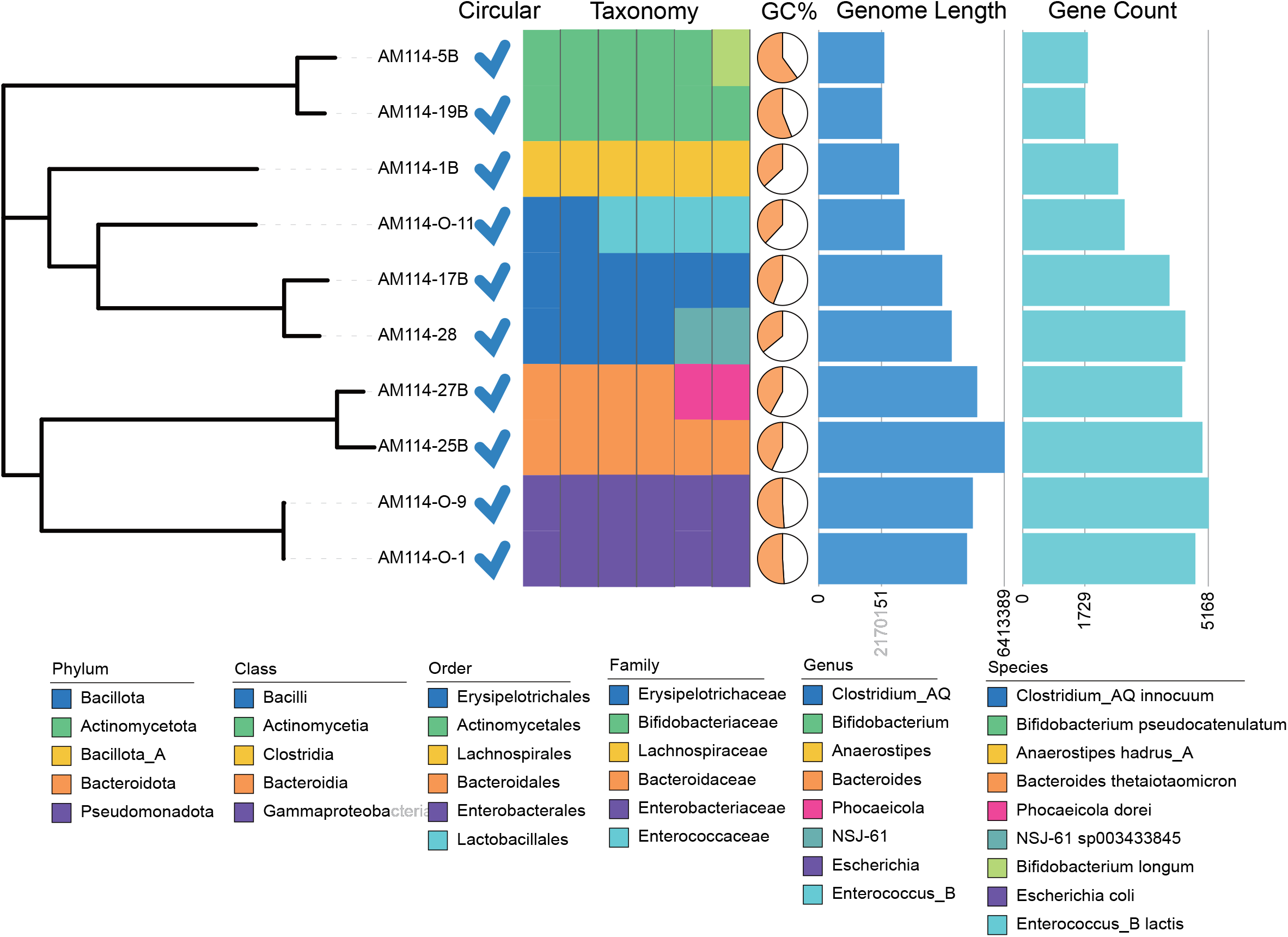
The phylogenetic tree of the 10 strains. All the genomes are circular. Different taxonomic levels and classification information are indicated with relevant colors. In the pie chart, the orange coverage represents the GC content. The length and number of genes for each genome are indicated by bar plots.

For these 10 test samples, we further analyzed the circular genomes by hybrid assembly. According to GTDB taxonomic annotation [19], they could be classified into 5 phyla, 5 classes, 6 orders, 6 families, 8 genera, and 9 species, which included two strains of *Escherichia coli* (**Fig. 2**). The GC content of these strains varied from 36% to 60%. The size of the genomes ranged from 2.17 Mbp to 6.41 Mbp, and the gene counts ranged from 1,729 to 5,168. These assemblies suggest that the hybrid assembly approach, integrating DNBSEQ short-read and CycloneSEQ long-read, capitalizes on the strengths of both long and short reads to assemble complete and accurate genome assemblies for common types of bacteria.

### Long reads restore multi-copy genes by filling gaps

Hybrid assembly effectively enhances genome completeness, and the improvements brought by long reads to the draft assembly are noteworthy. Evaluating these ten diverse genomes from the perspective of basic functional elements, the complete genome shows a significant increase in the number of CDS, rRNA, and tRNA coding genes compared to the draft (**Fig. 3A**). The increase in the number of rRNA is particularly notable, including 5S rRNA, 16S rRNA, and 23S rRNA. The draft can only assemble 2-7 rRNA genes, while the hybrid assembled genome can assemble 8-22 rRNA genes. 5S rRNA, 16S rRNA, and 23S rRNA often appear as a cluster close to 4,500 bp in length in the genome (5S rRNA: 68-111bp, 16S rRNA: 1519-1558bp, 23S rRNA: 2869-3056bp) (**Supplementary Fig. 4A,B**), and they present as multiple copies at different locations in the genome map (**Fig. 3B**), which is highly challenging for short reads assembly. In the draft genome assembled using short reads, there is often only one set of rRNA, while other regions containing rRNA become gaps.

**Figure 3.**
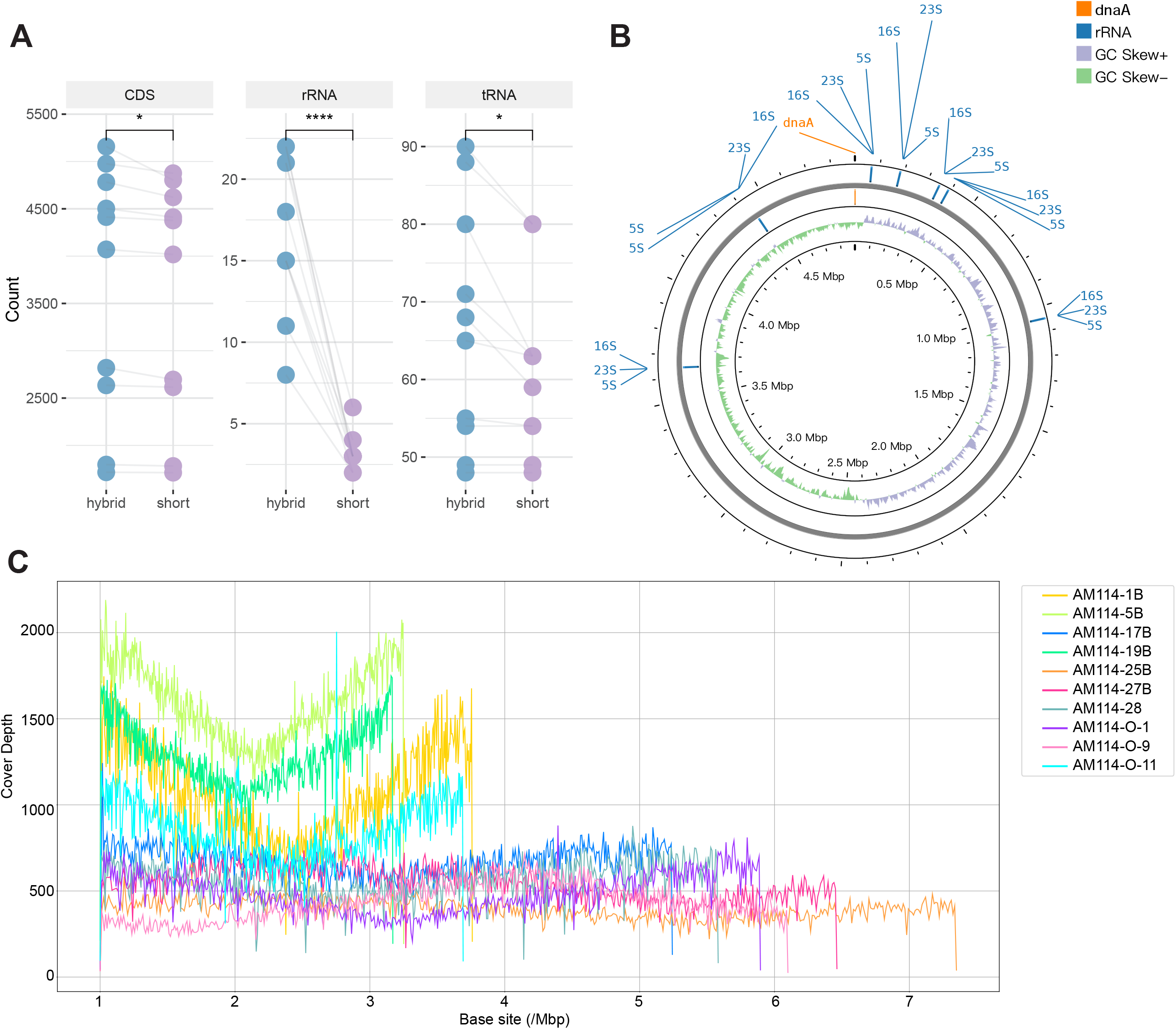
Comparison of complete genomes and short-read assemblies. **A**. The gene count annotated as coding sequences (CDS), ribosomal RNA (rRNA), and transfer RNA (tRNA). The complete genomes and short-read assemblies of the same strain were linked, and the paired values were compared using the Wilcoxon test. **B**. The circos plot of the chromosome genome of AM114-O-1, with all 22 rRNA positions indicated on the graph. **C**. Covering depth of each base position on the complete genome by short-read sequencing.

**Figure 4.**
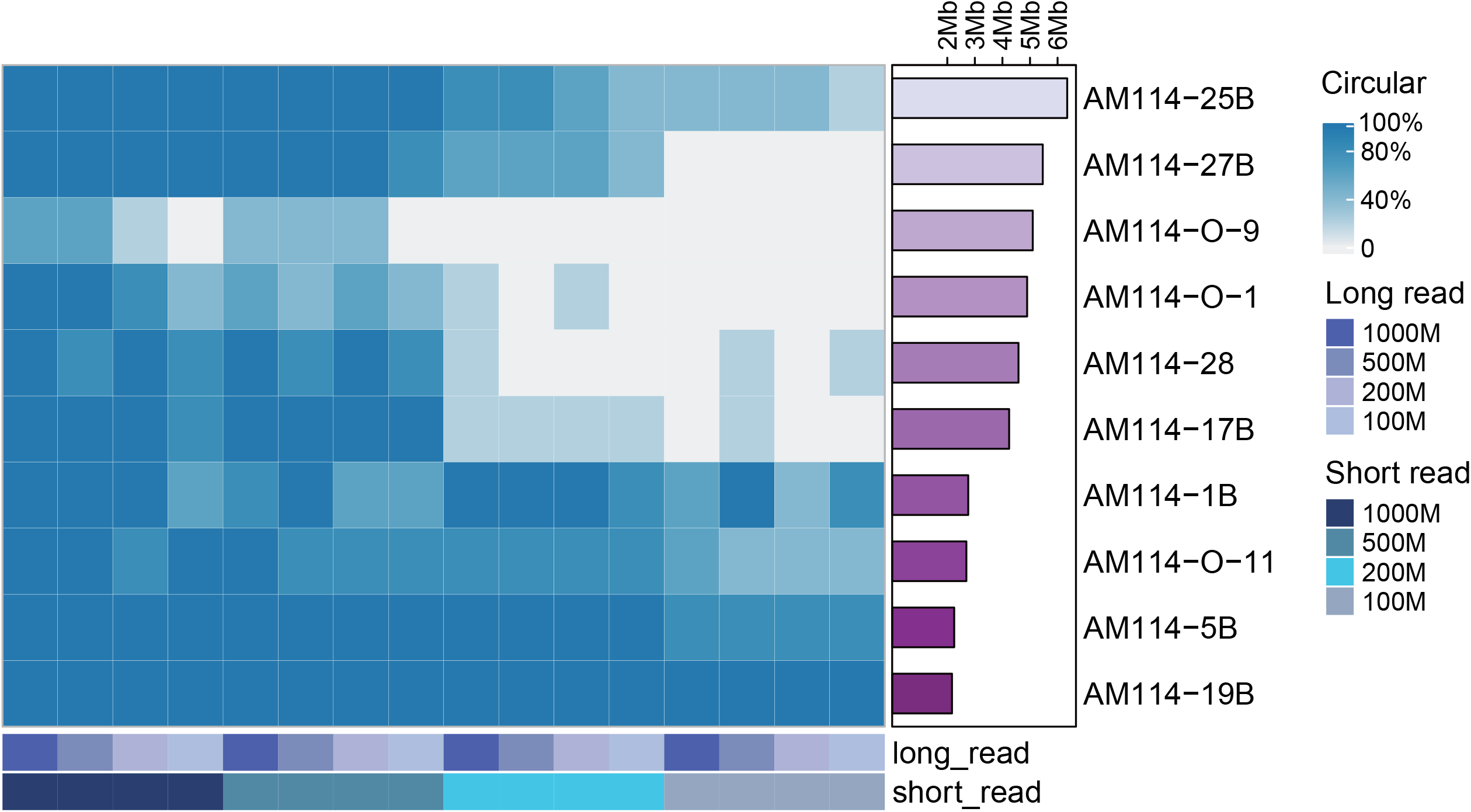
The proportion of circular genomes assembled under different data volumes. For each strain, 5 sets of data with volumes of 100Mb, 200Mb, 500Mb, and 1000Mb are randomly sampled for hybrid assembly. The proportion is calculated based on the number of circular chromosomes formed in the 5 genomes.

By mapping short reads to the complete genome, all the base sites of the AM114-1B, AM114-5B, and AM114-19B genomes have a coverage depth of more than 100X (**Figure 3C, Supplementary Table S3**). However, there are still gap regions in the short-read assemblies. In other complete genomes, there are continuous sites with depths fewer than 100X (**Supplementary Table S3**), indicating that both the lack of short-read sequencing depth and multicopy regions contribute to the gap regions in short-read assemblies. When long reads are mapped to the complete genome, there are reads longer than 5 kbp that can cover the gap regions in the draft, acting as a bridge and guiding genome assembly, thereby making the connections possible.

### Optimal data volumes for complete circular genome assembly

In order to determine the minimum data volume required for assembling complete circular genomes, we randomly generated 5 repetitions of the long-read and short-read sequencing data from 10 actual samples, with subset sizes of 100 Mbp, 200 Mbp, 500 Mbp, and 1000 Mbp each. This resulted in a total of 400 subsets (5 replicates * 4 subset sizes * 2 read types * 10 samples). Subsequently, we permuted these subsets from the same sample and assembled them.

When 1000 Mbp of short-read data was combined with either 1000 Mbp or 500 Mbp of long-read data, 9 out of 10 or 8 out of 10 samples respectively achieved 100% (5/5) assembly into circular genomes (**Fig. 4**). Remarkably, even with 200 Mbp or 100 Mbp of long-read data, respectively, 7 out of 10 or 5 out of 10 samples achieved complete assembly into circular genomes in all five replicates. Overall, when using 1000 Mbp of short-read data as a base and combining it with 100Mbp, 200Mbp, 500Mbp, 1000Mbp of long-read, the rates of achieving circular complete genomes were 76%, 88%, 94%, 96% respectively. This result suggests that for the assembly of bacterial genomes of common species and sizes, using 1000 Mbp of short-read data combined with more than 100 Mbp of long-read data greatly increases the likelihood of achieving a complete genome assembly.

Hybrid assemblies using 500 Mbp of short-read data were slightly inferior to those using 1000 Mbp of short-read data for the strains, but overall, they still achieved complete genome assembly at rates of 88%, 84%, 84%, and 74%, respectively. When the volume of short-read data is 200 Mbp, the rates of complete genome assembly are only around 50%; even when combined with 1000 Mbp of long-read data, only 3 out of 10 strains are fully assembled into circular genomes across all five replicates (**Fig. 4**). And when using only 100 Mbp of short-read data, the rates of complete genome drops markedly, resulting in an overall rate of only 34.5% (**Supplementary Table S4**). It is noteworthy that in assembly approaches reliant on short-read, the volume of short-read data is crucial for achieving a complete assembly. When there is an adequate volume of short-read data, just a few hundred Mbp of long-read data can suffice to produce complete assembly excellently.

### Impact of data volume on genome assembly accuracy

The integration of long-read data into the assembly process significantly enhances both the completeness and the rate of complete genome assembly. Concurrently, the accuracy of the assembled genomes remains an important consideration. Further, we utilized the value of indels and mismatches calculated by QUAST [20] to evaluate the accuracy of genomes assembled from subsets of varying sizes, using the hybrid assembly from the original full dataset as the reference.

With 1000 Mbp of short-read data, 76% of assemblies have mismatches below 1 bp per 100 kbp, and 97.5% have fewer than 10 bp per 100 kbp, with all assemblies exhibiting fewer than 10 indels per 100 kbp (**Fig. 5**). When the short-read data is reduced to 500 Mbp, there is a slight decline in performance compared to 1000 Mbp; however, 71% of assemblies still have mismatches below 1 bp per 100 kbp, 95% remain under 10 bp per 100 kbp, and all maintain fewer than 10 indels per 100 kbp, maintaining a high quality of assembly. In contrast, reducing the short-read data volume to 200 Mbp or 100 Mbp leads to a significant increase in the error rate for mismatches and indels per 100 kbp, with only 37% and 21% of assemblies, respectively, having mismatches under 1 bp per 100 kbp. Moreover, the average number of mismatches rises to 6.09 and 11.16 per 100 kbp. The above demonstrates that short-reads are particularly crucial for controlling the error rate.

**Figure 5.**
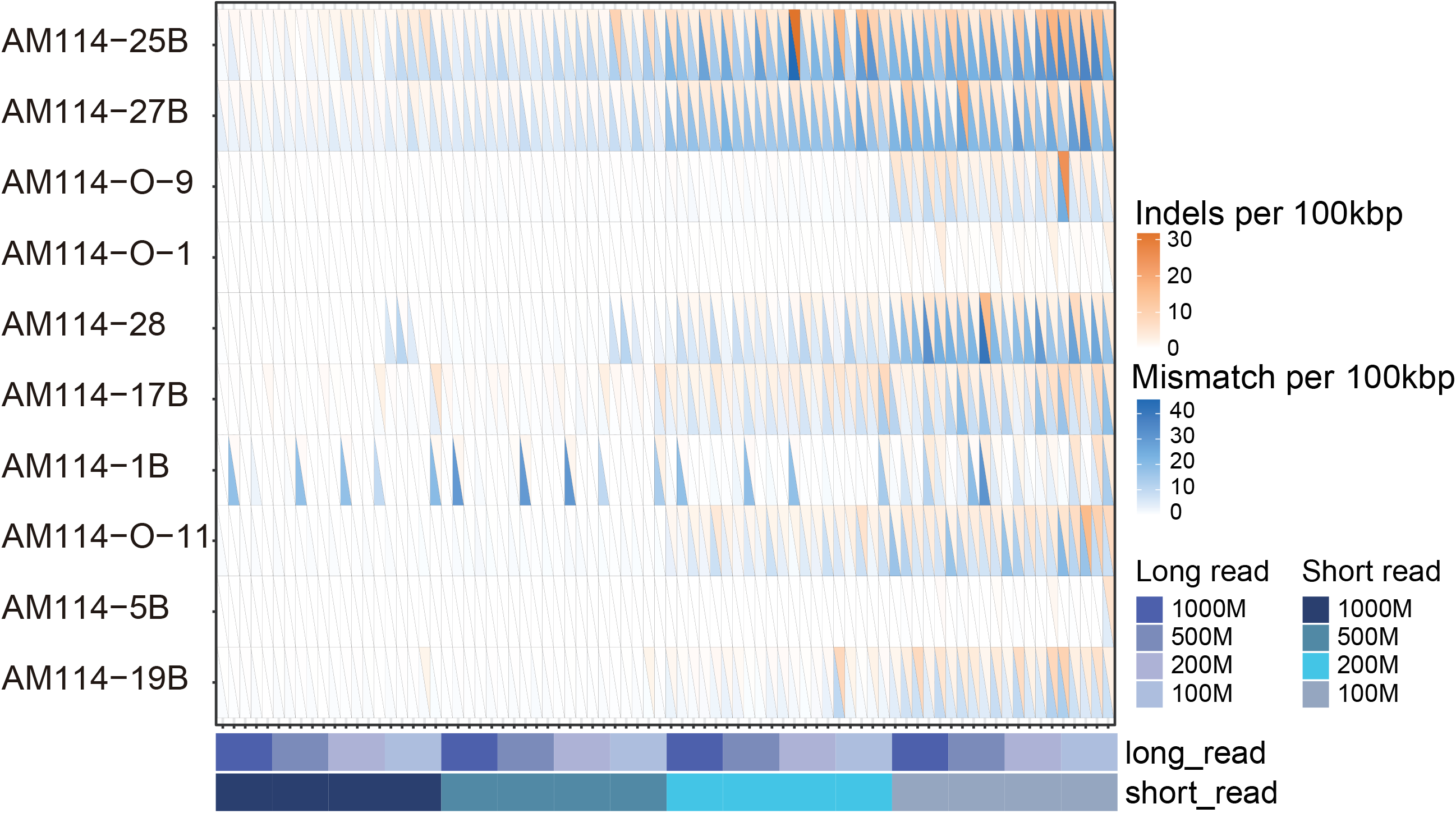
The error rate of assemblies under different data volumes. The genomes were the same as in Figure 4. Mismatches per 100kbp and indels per 100kbp are denoted by blue and orange colors, respectively, in the heatmap.

### Feasibility of assembly for microbial communities

Metagenomics is an important application in the field of microbiology. To evaluate the performance of CycloneSEQ in assembling mixed microbial communities, we used the Gut Microbiome Standard, which includes 18 bacterial strains, 2 fungal strains, and 1 archaeal strain, for assessment. To evaluate the performance of short-read assembly, long-read assembly, and hybrid-assembly, some common-used assembly methods were used for sequence assembly. And metagenome-assembled genomes (MAGs) were produced by binning from short-read assembly (metaSPAdes [21]), long-read assembly (metaFlye [18]), and hybrid-assembly (Unicycler, metaSPAdes, OPERA-MS [22]).

Under the same computing condition, hybrid-assembly methods consume more time than short-read assembly by metaSPAdes and long-read assembly by metaFlye (**Fig. 6A**). Unicycler was the most time-consuming method for hybrid-assembly, two-time more time consumed than metaSPAdes and OPERA-MS. Long-read assembly produced fewer MAGs than short-reads assembly and hybrid-assembly (**Fig. 6B**). Short reads played a key role in improve the number of MAG. Hybrid-assembly by Unicycler and metaSPAdes produced more single-contig MAGs (**Fig. 6C**). The completeness of MAGs from long-read assembly was lower than references and MAGs from short-read hybrid-assembly methods (**Fig. 6D, Supplementary Fig. 6C**). The contamination of MAGs from OPERA-MS (**Fig. 6E, Supplementary Fig. 6D**) and metaFlye (**Supplementary Fig. 6D**) were higher than references. N50 of MAGs produced by short-read assembly was shorter than MAGs from long-read and hybrid-assembly methods (**Fig. 6F, Supplementary Fig. 6E**). N50 of MAGs produced by OPERA-MS only slightly higher than N50 of MAGs by short-read assembly. In hybrid-assembly categories, metaSPAdes was the recommended tool for hybrid assembly of shot reads and long reads.

**Figure 6.**
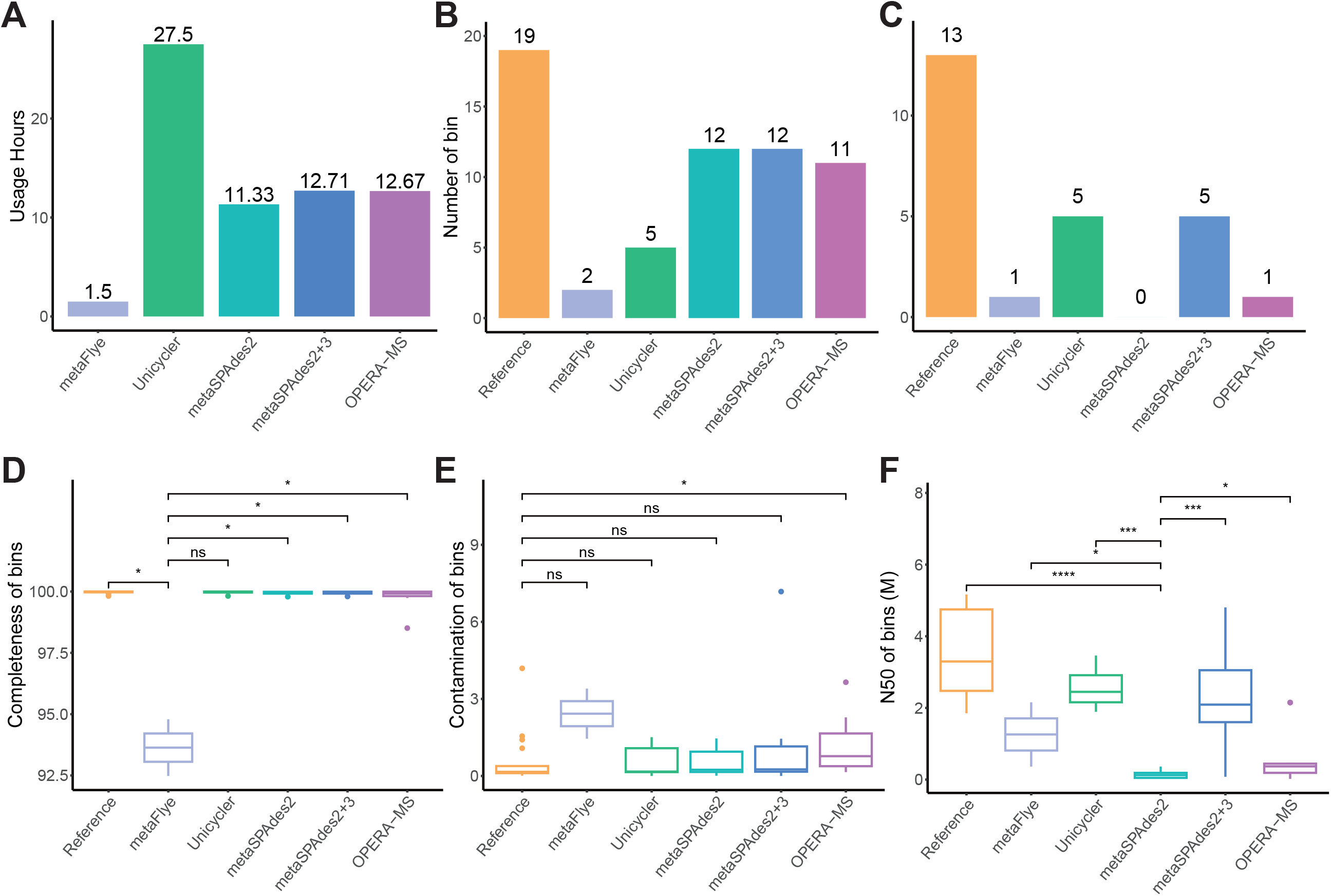
High-quality MAGs from short-read assembly, long-read assembly, and hybrid assembly for mock metagenomic sequences. **A**. The time consumption of sequence assembly. **B, C**. Number of MAGs and single-contig MAGs. **D-F**. Completeness, contamination, and N50 of MAGs. MAGs: completeness >= 90%; contamination <= 10%. Long-read assembly: metaFlye; short-read assembly: metaSPAdes2; hybrid assembly approach: metaSPAdes2+3 and OPERA-MS. *, p value < 0.05; **, p value < 0.01; ***, p value < 0.001; ****, p value < 0.0001; ns, p value > 0.05.

## Discussion

As a newly developed nanopore long-read sequencing platform, CycloneSEQ has demonstrated its sequencing performance and practical application in microbiology through this study. With an average read length of 11.7 kbp, comparable to other nanopore sequencing platforms, our analysis confirmed that this length is sufficient for assembling the circular bacterial genomes. However, CycloneSEQ has notable deficiencies in base quality. Therefore, integrating high-quality short reads by DNBSEQ is a promising solution. Ultimately, we achieved a high-quality assembly with only 0.08 mismatches and 0.15 indels per 100 kbp compared to the reference, this result may also be influenced by strain variations during cultivation.

We evaluated the performance of CycloneSEQ and DNBSEQ on 10 bacterial strains. Sequencing and assembling genomes using only short reads, only long reads, and a hybrid of both, we found that hybrid assemblies consistently produced high-quality circular genomes, including potential small circular genomes from bacteriophages or plasmids. Our tests indicate that for some common species with GC content ranging from 36% to 60% and genome lengths between 2.17 Mbp and 6.41 Mbp, the hybrid assembly approach can successfully produce circular genomes.The hybrid approach of integrating DNBSEQ short reads and CycloneSEQ long reads effectively combines their strengths, producing more complete and accurate genome assemblies than either method alone. This improvement is particularly evident in the increased number of CDS, rRNA, and tRNA coding genes in the complete genome compared to the draft. Due to limitations, we did not conduct sequencing tests on many more species or strains under more extreme conditions.

The ability to achieve high-quality assemblies with reduced long-read data volumes can make the hybrid assembly approach more cost-effective and accessible for various genomic research applications. This balance between data volume and assembly quality is crucial for optimizing resources in genomic studies. Using 1000 Mbp of short-read data combined with varying amounts of long-read data, we achieved high success rates in assembling complete circular genomes, with up to 96% success. Even with reduced long-read data of 500 Mbp, the success rates remained robust. In the meanwhile, the volume of short-read data significantly impacts assembly accuracy. With 1000 Mbp of short-read data, 76% of assemblies had fewer than 1 bp mismatches per 100 kbp, and 97.5% had fewer than 10 bp mismatches per 100 kbp. Reducing short-read data to 500 Mbp slightly decreased performance but maintained high accuracy. Further reduction to 200 Mbp or 100 Mbp led to a significant increase in error rates. These findings highlight that while long-read data is crucial for achieving complete genome assemblies, sufficient short-read data is essential for maintaining high accuracy. The hybrid assembly approach effectively balances these needs, making it the efficient method for bacterial genome assembly. In conclusion, the hybrid assembly approach, particularly with adequate short-read data, is the optimal method for bacterial genome assembly, balancing completeness, accuracy, and cost-effectiveness.

Real microbial samples have complex microbial compositions and biochemical components, making metagenomic sequencing and assembly more challenging for nanopore sequencing platforms. To effectively evaluate the feasibility of CycloneSEQ in sequencing mixed microbial communities, we used the Gut Microbiome Standard as a substitute for metagenomic samples. Overall, using the metaSPAdes tool for hybrid assembly of CycloneSEQ long reads and DNBSEQ short reads effectively combines the advantages of both short and long reads to produce complete and accurate genome assemblies. These findings report the utility of CycloneSEQ in metagenomics and highlight the advantages of hybrid assembly approaches. It is important to note that our study did not test real clinical samples, as the varying biochemical compositions between different samples could affect sequencing to different extents. This requires us to design more rigorous tests to achieve fair results. Future research should systematically test more real samples and focus on further optimizing the balance between short-read and long-read data to enhance assembly quality and efficiency. The CycloneSEQ long-read sequencing platform will facilitate these advancements in microbiome research.

## Methods

### Sample collection, DNA extraction, library construction, and sequencing

A fecal sample was collected from a healthy man, the collection was approved by the Institutional Review Board of BGI Ethical Clearance under number BGI-IRB 22112-T1. Sample was diluted and spread on agar culture mediums with anaerobic condition. We then picked 114 single colonies and transferred each of them to 2ml of liquid medium for further culture. 16S rDNA PCR was performed to identify the species, then we selected 10 diverse strains belonging to 9 species for further sequencing. The type strain ATCC-BAA-835 DNA was extracted using the Qiagen QIAamp DNA Mini Kit for long DNA fragments, while the test strain DNA was extracted using the Magen MagPure DNA Kit for high-throughput applications. The CycloneSEQ library preparation and sequencing followed the manufacturer’s guidelines. Each sample, containing 2 μg of input DNA (≥21 ng/μL), was diluted with nuclease-free water to 192 μL, then mixed with 14 μL each of DNA repair buffers 1 and 2, 12 μL of DNA repair enzyme 1, and 8 μL of DNA repair enzyme 2. The mixtures were incubated in a thermocycler at 20? for 10 minutes, 65? for 10 minutes, and held at 4?. After incubation, the mixtures were purified with 1.0x DNA clean beads and eluted with 240 μL of nuclease-free water. The end-repaired samples were then mixed with 10 μL of sequencing adaptors, 100 μL of 4x ligation buffer, 40 μL of DNA ligase, and 10 μL of nuclease-free water, and incubated at 25? for 30 minutes. The ligated products were purified again with 1.0x DNA clean beads, resuspended with long fragment wash buffer, and recovered into 42 μL of elution buffer. The libraries were quantified using a Qubit fluorometer and sequenced on the CycloneSEQ WuTong02 platform according to the protocol.

### Quality Control and Data Evaluation

Long-read data was filtered using NanoFilt [23] with parameters “-q 10 -l 1000” to retain reads longer than 1000 bp and with a quality score greater than Q10. Short-read data was processed using Fastp [24] with default parameters, except the length requirement was set to 50. The quality information of the data was evaluated using the tool seqtk [25], selecting the avgQ value from the ‘fqchk’ module as the average quality. The read lengths were extracted using a python script, and a density plot was generated based on this information.

### Short-read, long-read, and hybrid assembly of the isolated genome

Short-read assembly was performed using Unicycler [16] with only the short reads ‘-1’ and ‘-2’ as input, and the ‘–depth_filter’ set to 0.01 to remove low-depth contigs. For hybrid assembly, the same ‘depth_filter’ of 0.01 was used, with the addition of ‘-l’ long reads as input, while all other parameters were set to default. For long-read assembly, we used Flye [26] with the filtered long reads as input using the ‘–nano-hq’ option.

### Data splitting

Data splitting was performed using a custom Python script. Based on the required data volume, the script divided the data into FASTQ files of different sizes. It is important to note that here, Mb represents 1,000,000 bases, not the 1024-based system. Each read was treated as a unit, and the ‘random.sample()’ function was used for random selection. Reads were added one by one, and the total number of bases was calculated. When the addition of the last read met the required base count, the desired file was obtained. For paired short-reads, we assigned a sequence number to each read in the _1 and _2 files. Paired reads were then obtained by randomly selecting these sequence numbers.

### Completeness assessment and comparative evaluation of genome assemblies

The completeness of the genome was assessed using CheckM2 [27], while circularity was evaluated from the assembly results using Unicycler and Flye. We used QUAST [20] software for reference-based comparisons. To evaluate the assembly of the type strain, we used the genome ‘GCA_000020225.1’ from GenBank as the reference. For the evaluation of real samples, we used the hybrid assembly results from the full data set as the reference. Gene prediction and annotation were performed using Prokka [28], with coding sequences identified by Prodigal [29], and rRNA predicted using Barrnap.

### Read mapping to genome and depth calculation

Bowtie2 [30] was used to map the short reads to the complete genomes with the ‘–very-sensitive’ option. Samtools [31]was then used to convert the Bowtie2 output .bam file to the depth of each base site.

### Assembling, binning, annotation, and assessment for mock metagenomic data

SPAdes [21] (v3.15.5, -meta) was used for short-read assembly. SPAdes (v3.15.5, -meta), OPERA-MS [22] (v0.9.0), and Unicycler [16] (v0.5.0, -l) were used to hybrid assembly of short reads and long reads. Flye [26] (v2.9.3, --meta --nano-raw) was used for long reads assembling. 120Gb memory and 24 threads were prepared for sequence assembly. Metagenome-assembled genomes (MAGs) were constructed by Metawrap [32] (v1.3.2, -metabat2 -maxbin2 -concoct). Assembled genomes were annotated by GTDB-tk [19] (v2.3.2). Completeness and contamination of MAGs were assessed by CheckM2 (v1.0.1), and genomic quality assessment were conducted by Quast [20] (v5.2.0). ANIs between MAGs and reference genome of mock metagenome were calculated by FastANI [33] (v1.33). R software (v4.1.1) was used for data analysis and data visualization.

## Acknowledgements

This work was supported by grants from the Shenzhen Municipal Government of China (No. XMHT20220104017, CXB201108250097A). We also thank the colleagues at BGI-Shenzhen for sample collection, and discussions, and China National GeneBank (CNGB) Shenzhen for DNA extraction, library construction, and sequencing.

## Author contributions

Conceived and designed the study: Y.Z., L.X., X.X., H.L., C.L., X.J., W.Z.. Performed the analysis: H.L., M.W., T.H., H.W., W.H., Y.W., L.X., Y.J., R.G. Contributed reagents/materials/analysis tools: J.C., F.G., T.Z., B.W., X.J., X.X., J.W. Wrote the paper: Y.Z., Z.J., H.L., Y.M., J.Z. Supervised the work: L.X., X.X., Y.Z. All authors commented on the manuscript.

## Data availability

The data that support the findings of this study have been deposited into CNGB Sequence Archive (CNSA)[34] of China National GeneBank DataBase (CNGBdb)[35] with accession number CNP0006129.

